# Control of *φ* C31 integrase-mediated site-specific recombination by protein trans splicing

**DOI:** 10.1101/540872

**Authors:** Femi J. Olorunniji, Makeba Lawson-Williams, Arlene L. McPherson, Jane E. Paget, W. Marshall Stark, Susan J. Rosser

## Abstract

Serine integrases are emerging as core tools in synthetic biology and have applications in biotechnology and genome engineering. We have designed a split-intein serine integrase-based system for rapid regulation of site-specific recombination events *in vivo.* The *φC31* integrase was split into two extein domains, and intein sequences *(Npu* DnaE^N^ and *Ssp* DnaE^C^) were attached to the two termini to be fused. Expression of these two components followed by post-translational protein *trans-splicing* in *E. coli* generated a fully functional *φC31* integrase. Protein splicing is necessary for recombination activity; no activity was observed when the *φ*C31 integrase N-and C-terminal extein domains without the intein sequences were co-expressed, nor when a key intein catalytic residue was mutated. As a proof of principle, we used a bistable switch based on an invertible promoter reporter system to demonstrate a potential application of the split intein-regulated site-specific recombination system. We used *araC* and *tet* inducible promoters to regulate the expression of the two parts of the split recombinase. Inversion of a DNA segment containing a constitutive promoter, catalyzed by *trans*-spliced integrase, switches between RFP and GFP expression only when both inducible promoters are ON. We used the same split inteins to regulate the reconstitution of a split integrase-RDF fusion that efficiently catalyzed the reverse *attR* x *attL* recombination, demonstrating that our split-intein regulated recombination system can function as a reversible AND gate in which the forward reaction is catalyzed by the integrase, and the reverse reaction by the integrase-RDF fusion. The split-intein integrase is a potentially versatile, regulatable component for building synthetic genetic circuits and devices.

## INTRODUCTION

It has recently become possible to create computational and memory systems in cells^1,2,3^ allowing us to foresee many new ways to enhance the applications of living organisms^4,5^. Engineered cells could act as powerful biosensors with applications in health, environmental, and industrial processes. As well as sensing components, it is necessary to process the received information and express it as outputs in the form of specific biological responses^3,6^. However, the genetic switches and logic gates that have been constructed to date are based on a limited repertoire of biological component types^7^, and there is a need for new systems that can be used to implement more elaborate and robust devices.

DNA site-specific recombination has been much exploited for rapid DNA assembly, and to build genetic switches and memory devices^4,8,9,10^. In a typical module, two recombination sites flank a promoter sequence. Expression of the recombinase promotes inversion of the orientation of the promoter sequence, thus switching between expression of two genes which are divergently transcribed from the module (Figure 1A). One group of site-specific recombinases known as the serine integrases is especially suited for the construction of switching devices, particularly because these enzymes promote very efficient and highly directional recombination^11^. For such modules to be useful, fully integrated components of the cell, activity of the recombinase must be tightly regulated, so that switching occurs only when other cellular conditions are fulfilled. Here we demonstrate a powerful new approach to regulation of serine integrase activity, in which the enzyme itself is assembled by intein-mediated fusion of two precursor components.

**Figure 1:**
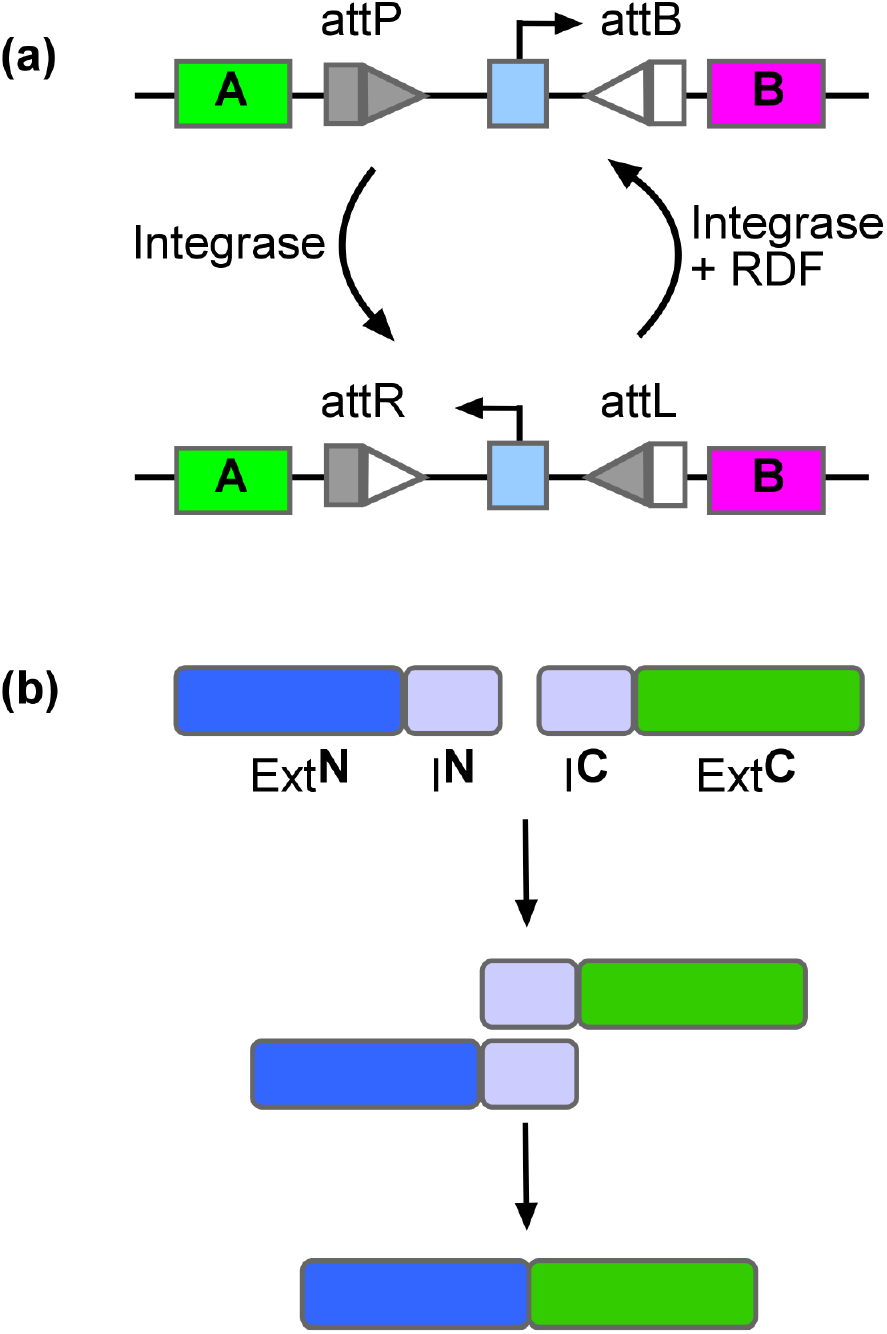
Building blocks for split intein-mediated trans-splicing of *φ* C31 integrase. **(A)** Serine integrase-catalysed inversion of the orientation of a promoter sequence functions as a toggle switch for the expression of two genes. Integrase catalysed recombination between *attP* and *attB* sites (arranged in inverted orientation) results in switching the direction of the promoter, and hence changes the gene expression status of the module. Recombination of *attR* and *attL* sites by integrase and the recombination directionality factor (RDF), restores the module to its original state. **(B)** Split-intein catalyzed protein trans-splicing. Non-covalent association of the N-terminal intein (**I^N^**) and the C-terminal intein (**I^C^**) is followed by a trans-splicing reaction that covalently joins the N-terminal extein (**E^N^**) and C-terminal extein (**E^C^**).

Inteins are naturally occurring autocatalytic systems that catalyse protein splicing reactions to generate active proteins from precursor polypeptides^12^. Synthetic “split inteins” have been developed to carry out protein splicing *in trans,* covalently joining two proteins together for a wide range of biotechnological applications^13,14^ (Figure 1B). For example, Schaerli^15^*et al.* split T7 polymerase into two parts and fused each part to split-intein sequences so that the two parts were covalently joined together by *trans*-splicing. Expression of each of the two individual parts was placed under the control of an inducible promoter to allow conditional expression of integrase activity.

A critical constraint on the construction of split-intein logic gates is the need to introduce an intein nucleophilic residue (typically Cys, Ser, or Thr) by mutation of the sequence of the target protein adjacent to the junction between the two ‘exteins’. Such changes can potentially lead to reduction or loss of enzyme activity. However, inteins have been engineered that tolerate variations of these flanking residues, thereby minimizing the number of changes that need to be made in the target protein^16,17^.

Conditional site-specific recombinase activation by assembly of split protein fragments has been achieved; for example, for the popularly used tyrosine recombinase Cre^18–22^. Wang *et al.* ^22^ used a Cre split-intein system to reconstitute functional recombinase in transgenic mice. However, there are no reports to date of analogous systems using split serine integrases. The transposase TnpX (distantly related to the serine integrases) was split into two parts and some DNA-binding activity was reconstituted when both parts were present, but no recombination (transposition) activity was observed^23^.

With the increasing importance of serine integrases as tools in synthetic biology, methods for signal-induced post-translational regulation of integrase activity are becoming very desirable. Here, we report post-translational activation of site-specific recombination by reconstitution of functional *φ*C31 integrase using split intein-catalysed reactions.

## RESULTS AND DISCUSSION

### Design of split intein serine integrase

We designed our split-intein system for conditional expression of *φ*C31 integrase, a prototype serine integrase, based on previously reported systems^15^. Since the natural DnaE split inteins used in this work require an invariant active site cysteine to remain in the spliced product protein^24–26^, their use for split integrase reconstitution depends upon the identification of a short region of the protein sequence where insertion of a cysteine does not disrupt activity. In addition, for *trans*-splicing to be essential, the two extein components of the spliced integrase must not associate non-covalently to reconstitute a functional enzyme. We started by analysing the domain structure of serine integrases to determine where to split the protein, since a functional split intein could require introduction of mutations that would remain in the spliced protein product, if suitably placed natural Cys, Ser, or Thr residues were unavailable. Based on sequence alignments of serine integrases (Figure 2A) and published crystal structures^27^, we split *φ*C31 integrase at the non-conserved region of the recombinase domain between the β9 and αI domains; this region includes a 10-12 residue loop which is absent in some related serine integrases^28^ (Figure 2A). We therefore predicted that introduction of the required intein nucleophilic residue and any flanking residues would not have a deleterious effect on integrase activity. Furthermore, Lucet *et al*^23^ found that when the related large serine recombinase TnpX (from *Clostridium perfringens)* was split between the β9 and αI domains, the two fragments complemented each other to restore DNA binding (but not recombination) activity.

**Figure 2:**
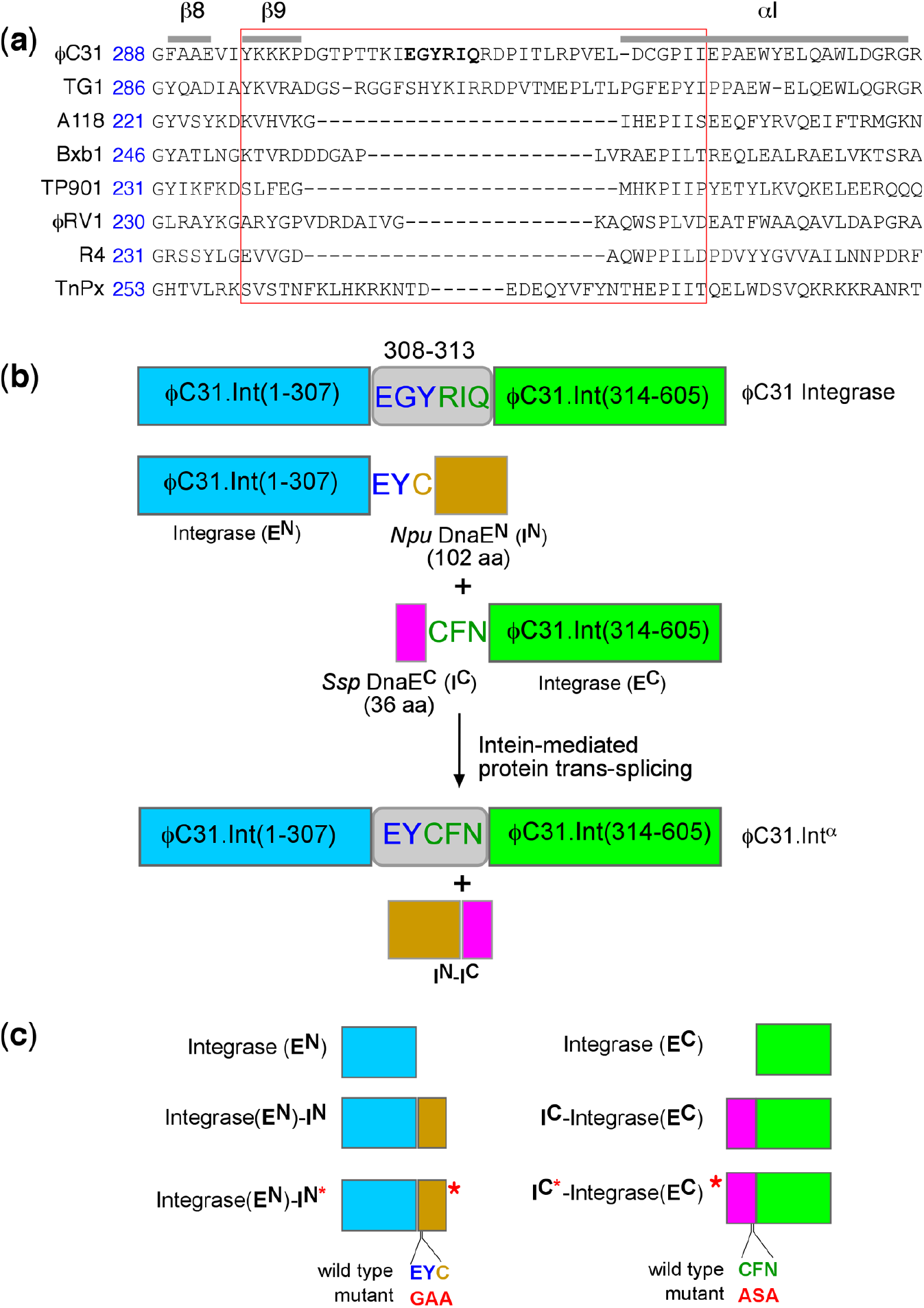
Design of split *φ*C31 integrase. **(A)** Sequence alignment of *φ*C31 integrase with related members of the ‘large serine recombinase’ protein family based on published structures and similar alignments^27^. Amino acid sequences in the loop region where *φ*C31 integrase is split are highlighted with a red box. **(B)** Structures of the split integrase-intein constructs. Changes made to the integrase sequence at the junctions where the fragments are fused to the split inteins are shown. The functional domains are shown as rectangular boxes: Integrase-**E^N^** (blue), *Npu* Dna**E^N^** (orange), *Ssp* Dna**E^C^** (pink), and Integrase-**E^C^** (green). Intein-catalysed protein *trans*-splicing generates an active version of the integrase in which residues at positions 308-312 of wild-type integrase (EGYRIQ) are replaced with EYCFN. The *trans*-spliced integrase is thus one amino acid shorter than the native integrase. **(C)** Variants of the split *φ*C31 integrase designed to probe the requirements for reconstitution of integrase activity. (i) Integrase-**E^N^**-**I^N^**; Integrase-**E^N^** (residues 1-307) fused to the 102-amino acid residue *Npu* Dna**E^N^** (**I^N^**) using the two-residue EY and the N-terminal Cys residue from *Npu* Dna**E^N^**. (ii) Integrase-**E^n^**; *φ*C31 integrase (1-307). (iii) Integrase-**E^N^**-**I^N^**\*; *φ*C31 integrase (1-307) fused to *Npu* Dna**E^N^** as described in (i) but with the EY residue at the splice site changed to GA to inactivate splicing activity. (iv) **I^C^**-Integrase-**E^C^**; *φ*C31 integrase C-extein (residues 314-605) fused directly to the *Ssp* Dna**E**^C^ (**I**^C^). *Ssp* Dna**E**^C^ nucleophilic cysteine residue and the flanking residues required for splicing are highlighted (CFN). (v) Integrase-**E**^C^; *φ*C31 integrase C-extein (residues 314-605). (vi) I^C^*-Integrase-**E**^C^; *φ*C31 integrase C-extein fused to *Ssp* Dna**E**^C^ as described in (i) but with the CFN residue at the splice site changed to ASA to inactivate splicing activity.

We then attached two well-characterised split-intein components to the *φ*C31 integrase sequences; *Npu* DnaE^**N**^, **I^N^** (102 amino acids) from *Nostoc punctiforme* DnaE, and *Ssp* DnaE^C^, I^C^ (36 amino acids) from *Synechocystis sp.* DnaE. The *Npu* DnaE^N^ variant contains a L22S change and the *Ssp* DnaE^C^ variant has a P21R mutation^24^. This pair was chosen based on previous reports of their optimal activities at 37 °C in *E. coli* ^15,24,29^. *Npu* DnaE^**N**^ (**I^N^**) was fused to the C-terminus of *φ*C31 Integrase (Integrase-**E^N^**), and *Ssp* Dna**E^C^** (**I^C^**) was fused to the N-terminus of *φ*C31 Integrase (Integrase-**E^C^**) (Figure 2B). Also, residues 308-310 of Integrase-**E^N^** were changed from EGY to EY, since these sequences exist naturally at the extein-intein junction of *Npu* DnaE^N^, and Integrase-**E^N^** residues 311-313 were changed from RIQ to CFN, sequences found at the intein-extein junction of *Ssp* DnaE^C^^24,29^. It is known that these flanking residues are involved in enhancing the splicing efficiencies of this pair of split inteins^25,29^. In the reconstituted *trans*-spliced integrase (φC31.Int^**α**^), the natural 6-residue sequence at positions 308 to 313, EGYRIQ, is replaced with the 5-residue sequence EYCFN. We made these changes to maximize the efficiency of the splicing reaction in order to optimize activity in *E. coli.* Hence the reconstituted *trans*-spliced integrase, *φ*C31.Int^**α**^ is one amino acid residue shorter than the wild-type *φ*C31 integrase. Since these changes are in a non-conserved region (see above), we predicted that they would not substantially affect recombination activity.

### Intein-mediated reconstitution of functional *φ* C31 integrase

To assay *in vivo* recombination activity of our split-intein *φ*C31 integrase, we used a well-characterized colour-based *galk* assay^30–33^. The *attP* and *attB* recombination sites (Supporting Information, Table S1) on the substrate plasmid p*φ*C31-delPB are in a direct repeat orientation, so that recombination between them causes deletion of the *galK* gene. Pale-coloured *(galK-)* colonies on the indicator plates indicate recombination proficiency, whereas red *(galK+)* colonies indicate incomplete or zero recombination. The coding sequences for Integrase-**E^N^-I^N^** and **I^C^**-Integrase-**E^C^** (and also appropriate control proteins; see Figure 3C) were cloned into separate low-level expression vectors^30^, as illustrated in Figure 3B. An *E. coli* strain containing the test substrate was transformed with these plasmids, and recombination activity was assessed by colony colour and by gel electrophoresis analysis of plasmid DNA recovered from the cells (Figure 3C).

**Figure 3:**
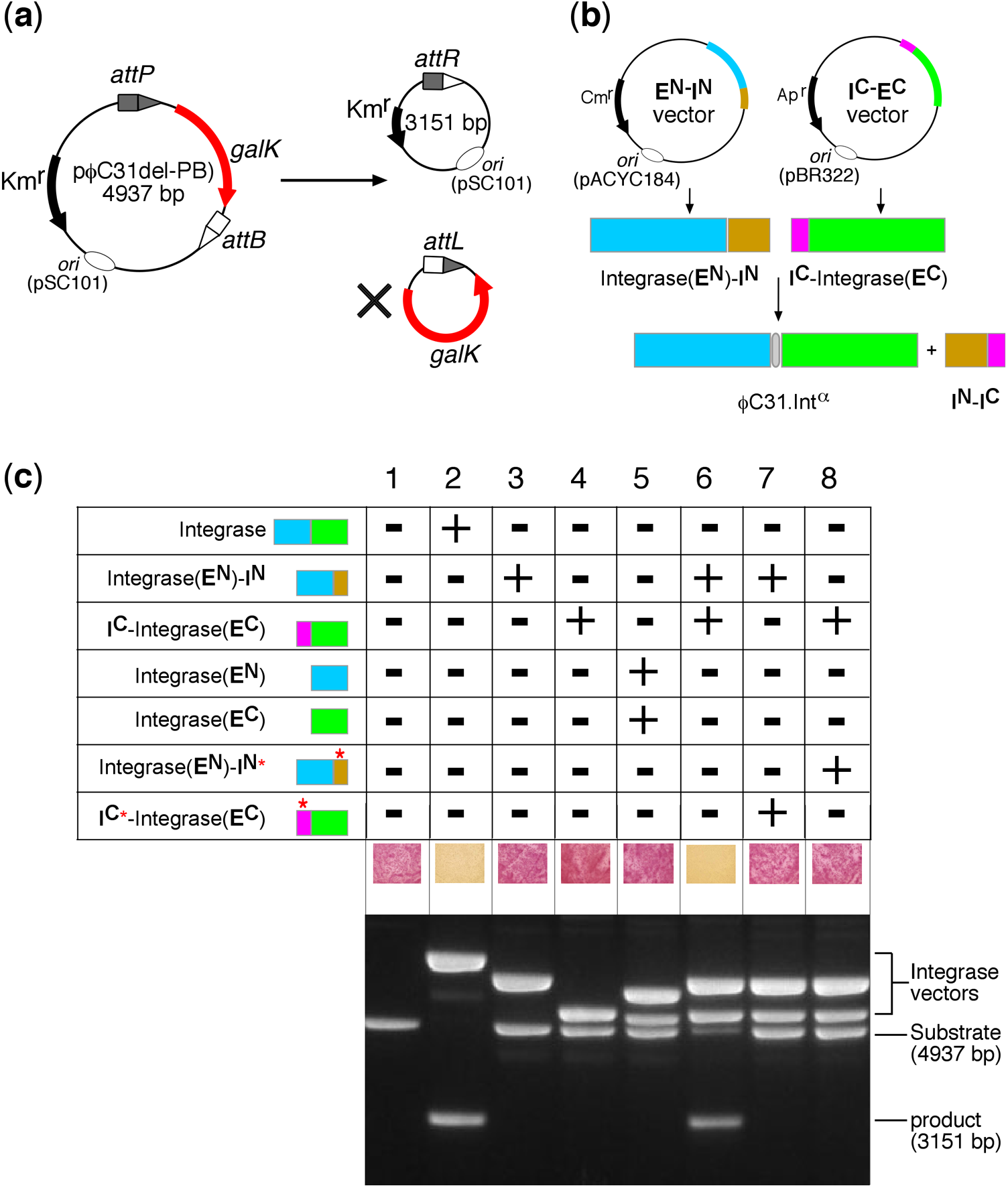
*In vivo* recombination activity of reconstituted split-intein *φ*C31 integrase. **(A)** *In vivo* recombination assay using a deletion substrate (p*φ*C31-delPB) carrying the *galK* gene^30^. The recombination product circle carrying the *galK* DNA has no origin of replication, and is lost during subsequent cell divisions. Inability of the cells to metabolize galactose leads to a change in colony colour on MacConkey galactose medium. (B) Constitutive expression of precursor polypeptides. The open reading frame of Integrase-**E^N^**-**I^N^** was expressed from a pACYC184-based vector, and **I**^C^-Integrase-**E**^C^ was expressed from a pBR322-based vector^30^. Expression of the two polypeptides followed by intein-catalysed protein trans-splicing reconstitutes *φ*C31.1nt^α^. (C) Assay of recombination activities *(galK* colour assay) of integrase constructs on *attP × attB* deletion substrates (p*φ*C31-delPB). DNA analysis of recombination activities of integrase constructs on *attP* × *attB* deletion substrates (p*φ*C31-delPB). In these assays, cells containing the substrate plasmid (p*φ*C31-delPB) were transformed with the expression vectors indicated and grown for 20 hours in selection media. For analysis of *in vivo* recombination products, plasmid DNA was recovered from cells^30^ and separated by means of 1.2% agarose gel electrophoresis. (1) Substrate only (blank control); (2) FEM141: *φ*C31 Integrase; (3) FEM136: Integrase-**E^N^**-**I^N^**; (4) FEM137: I^C^-Integrase-**E**^C^; (5) FEM155 (Integrase-**E^N^**) and FEM157 (Integrase-**E**^C^); (6) FEM136 (Integrase-**E^N^**-**I^N^**) and FEM137 (**I**^C^-Integrase-**E**^C^); (7) FEM136 (Integrase-**E^N^**-**I^N^**) and FEM162 (**I**^C^*-Integrase-**E**^C^); (8) FEM137 (**I**^C^-Integrase-**E**^C^) and FEM161 (Integrase-**E^N^**-**I^N^**\*).

Neither Integrase-**E^N^-I^N^** (Figure 3C, lane 3) nor **I^C^**-Integrase-**E^C^** (Figure 3C, lane 4) on their own were able to catalyse *attP* x *attB* recombination. Furthermore, co-expression of the N-extein (Integrase-**E^N^**) and C-extein (Integrase-**E^C^**) components of the integrase (lane 5) gave no recombination. This important control shows that the intein-less precursor proteins Integrase-**E^N^** and Integrase-**E^C^** do not complement each other by non-covalent association to give recombination activity. This contrasts with reported split tyrosine recombinases, the components of which associate to reconstitute recombination activity without protein splicing^18–20^. When the intein-tagged integrase exteins Integrase-**E^N^-I^N^** and **I^C^**-Integrase-**E^C^** were co-expressed, reconstitution of integrase recombination activity was observed (lane 6). The DNA analysis shown in the lower panel of Figure 3C suggests that recombination by the *trans*-spliced integrase (*φ*C31.Int^**α**^) has proceeded to over 80%.

Our results show that the changes from the wild-type integrase sequence that had to be introduced at the splice site are compatible with recombination activity. As many other serine integrases have similarly non-conserved, variable lengths of amino acid sequence in the region of the protein between β9 and αI (Figure 2A), we predict that these enzymes could also be engineered to create active split-intein systems.

### *Trans-splicing* is required for integrase recombination activity

The results shown in Figure 3C show that the *Npu* DnaE and *Ssp* DnaE intein moieties are required for reconstitution of recombination activity, but these experiments do not unambiguously establish the requirement for *trans*-splicing. Some split inteins are known to associate tightly via non-covalent interactions^34^, and this can lead to reconstitution of the split protein activity, without splicing^35^. To test whether this applies in our split integrase system, we mutated the nucleophilic cysteine residue and rate-enhancing flanking residues at the active sites of the two intein moieties^24^ to render them catalytically inactive. We changed the junction residues “EY” in the Integrase-**E^N^** moiety to “GA” and the Cys residue in the *Npu* DnaE^N^ moiety of Integrase-**E^N^**-**I^N^** to “A” to give the mutated version Integrase-**E^N^**-**I^N^**\* (Figure 2C). Similarly, the residues “CFN” in the *Ssp* Dna**E**^C^ moiety of **I**^C^-Integrase-E^C^ were changed to “ASA”, to derive the mutated version **I**^C^*-Integrase-**E**^C^ (Figure 2C). No recombination activity was observed when active Integrase-**E^N^**-**I^N^** was co-expressed with inactive **I**^C^*-Integrase-**E**^C^ (Figure 3C, lane 7), nor when active I^C^-Integrase-**E**^C^ was co-expressed with inactive Integrase-**E^N^**-**I^N^**\* (Figure 3C, lane 8), showing that reconstitution of recombination activity requires the catalytic activities of both split intein fragments. In this aspect, serine integrases differ from integrases of the tyrosine recombinase family where non-covalent association is sufficient to allow reconstitution of activity from split protein fragments^18–20^.

The *attP* x *attB in vivo* deletion reactions were slower in cultures where recombination required post-translational *trans*-splicing reactions to reconstitute functional integrase polypeptides, when compared to recombination by native integrase (see Supporting Information, Figure S1). It is likely that the rate of *trans*-splicing between *Npu* DnaE^N^ and *Ssp* DnaE^C^ is significantly slower *in vivo* in contrast to the fast *in vitro* splicing reaction rate^29^.

The post-translational control system described in this report is tightly regulated because significant expression of both extein components will be necessary to generate enough spliced integrase for any recombination reaction to happen. This offers an advantage over regulation at the transcriptional level, where ‘leaky’ promoters often result in unreliable output signals. The faster kinetics of intein-catalysed protein *trans*-splicing would enable rapid control of gene expression, in contrast to the slower method of conditional gene expression by means of transcriptional control.

### Split intein-regulated integrase-catalysed inversion system

To demonstrate the potential application of our split intein *φ*C31 integrase in conditional expression of recombination activity, we designed an invertible genetic system based on an inversion susbstrate plasmid, p*φ*C31-invPB. Recombination between *attP* and *attB* sites in p*φ*C31-invPB inverts the orientation of a constitutive promoter sequence (Biobrick J23104), thereby switching expression from RFP to GFP (Figure 4A). The *E. coli* strain DS941/Z1/p*φ*C31-invPB was co-transformed with two vectors, each expressing one of the two split-intein integrase fragments. Expression of Integrase-**E^N^**-**I^N^** was placed under the control of the *araC* promoter, and expression of **I**^C^-Integrase-**E**^C^ was placed under the control of the Ptet promoter (Figure 4B). Co-expression of the split-intein fragments (and thus recombination) is dependent on the presence of both of the inducers arabinose and anhydrotetracycline (aTc) (Figure 4C). No recombination (GFP expression) was observed unless both split-intein components were expressed (when aTC and arabinose were added to the growth medium; column IV.

**Figure 4:**
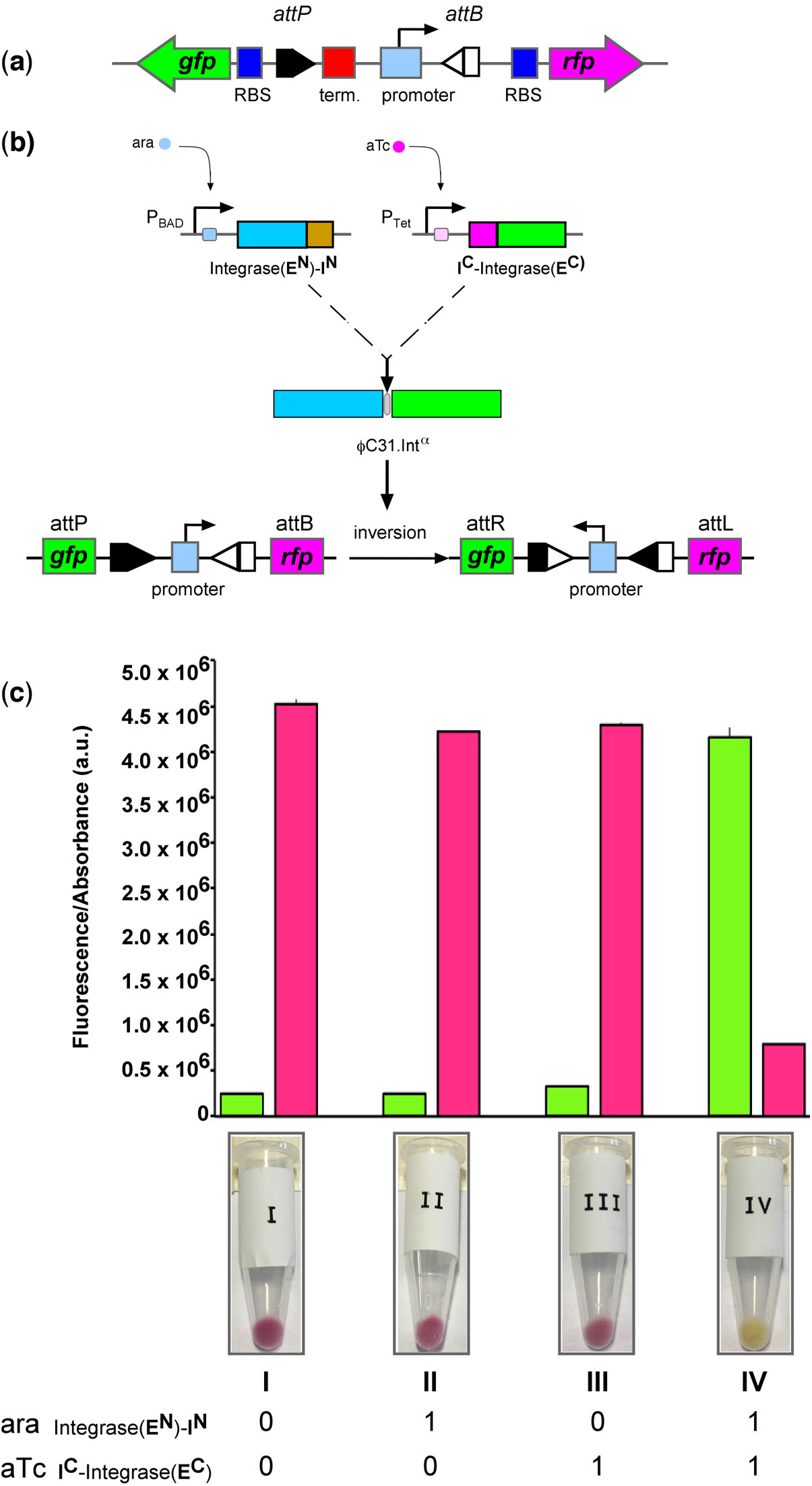
Controlling the function of an invertible promoter system using split intein-regulated *φ* C31 integrase activity. **(A)** Design of a recombinase-operated switch using an invertible promoter reporter system. The constitutive promoter sequence (cyan rectangle; Biobrick J23104) is flanked by *φ*C31 integrase *attP* and *attB* sites (grey arrows) that are arranged in inverted orientation. In its default state, the promoter constitutively drives the expression of a red fluorescent protein (*rfp*) gene (pink arrow). The T1 terminator sequence (red squares) located immediately upstream of the promoter prevents transcriptional read-through to the green fluorescent protein (gfp) gene (green arrow). Upon integrase-catalysed site-specific recombination, the orientation of the promoter is reversed to allow the expression of GFP, and block RFP production. A Biobrick ribosomal binding site, RBS, (RBS_B0034, blue rectangles), is positioned 5’ of the *rfp* and *gfp* genes to drive optimal translation of the synthesized mRNAs. **(B)** Conditional expression of *φ*C31 integrase fragments and split intein-mediated reconstitution of activity. The two split-intein integrase fragments are expressed from two inducible promoter vectors. Arabinose (ara) induces expression of Integrase-**E^N^**-**I^N^** under the control of the pBAD promoter (pFEM149), while anhydrotetracycline (aTC) induces expression of **I**^C^-Integrase-**E**^C^ under the control of the pTet promoter (pFEM148). Post-translational trans-splicing of the split-intein integrase fragments generates the functional reconstituted integrase (*φ*C31.Int^α^). Catalysis of *attP*x *attB* inversion by *φ*C31.Int^α^ results in reversal of the orientation of the promoter. **(C)** Validation of the split-intein *φ*C31 integrase recombinational AND-gate. Fluorescence measurements indicating the expression of GFP and RFP in cells induced with aTc and/or ara. Cells were grown at 37 °C for 24 hours in the presence of aTC (0.1 μg/ml) and ara (0.2%). Glucose was added to 0.4% concentration to treatments A and C to turn off expression of the arabinose promoter. Each bar represents mean and standard deviation of four determinations. Below the bar chart are images of Eppendorf tubes showing the pellet obtained after centrifugation of 5 ml of bacterial culture.

The split-intein regulated serine integrase can be deployed as an effective tool for transient inversion of genetic regulatory modules (Figure 4). The ability to reconstitute integrase activity from inactive protein fragments using split inteins could allow precise control of site-specific recombination activity, either by using promoters that respond to different chemical signals or by specific activation of the intein splicing reaction. In this work we have demonstrated a split-integrase system that could be engineered to function as components of a logic gate. This could be achieved by means of light-sensitive protein domains that regulate intein splicing^36,37^ or small molecule ligands that act directly on the split inteins^38^ to control the protein-protein association step that precedes protein *trans*-splicing. Others have demonstrated the use of pH changes^39^ and temperature^40^ as tools for regulating intein functions. These technologies can enhance capacity for building orthogonal logic gate components for conditional gene expression regulation and genome engineering applications, and add to the existing tools for programmable cellular functions^41,42^.

### Intein-mediated assembly of integrase-RDF fusion recombinase and catalysis of *attR* x *attL* recombination

We recently showed that recombination between *attR* x *attL* sites can be catalysed efficiently by artificial proteins in which the recombination directionality factor (RDF) is fused to the integrase using a short peptide linker. The fusion recombinase *φ*C31.integrase-gp3 catalysed *attR* x *attL* recombination efficiently^30^ (Figure 5A). This system allows more predictable regulation of integrase-catalysed conversion of *attP/attB* to *attR/attL* and vice versa, with potential applications in building genetic switches and recombinase-based counters.

**Figure 5:**
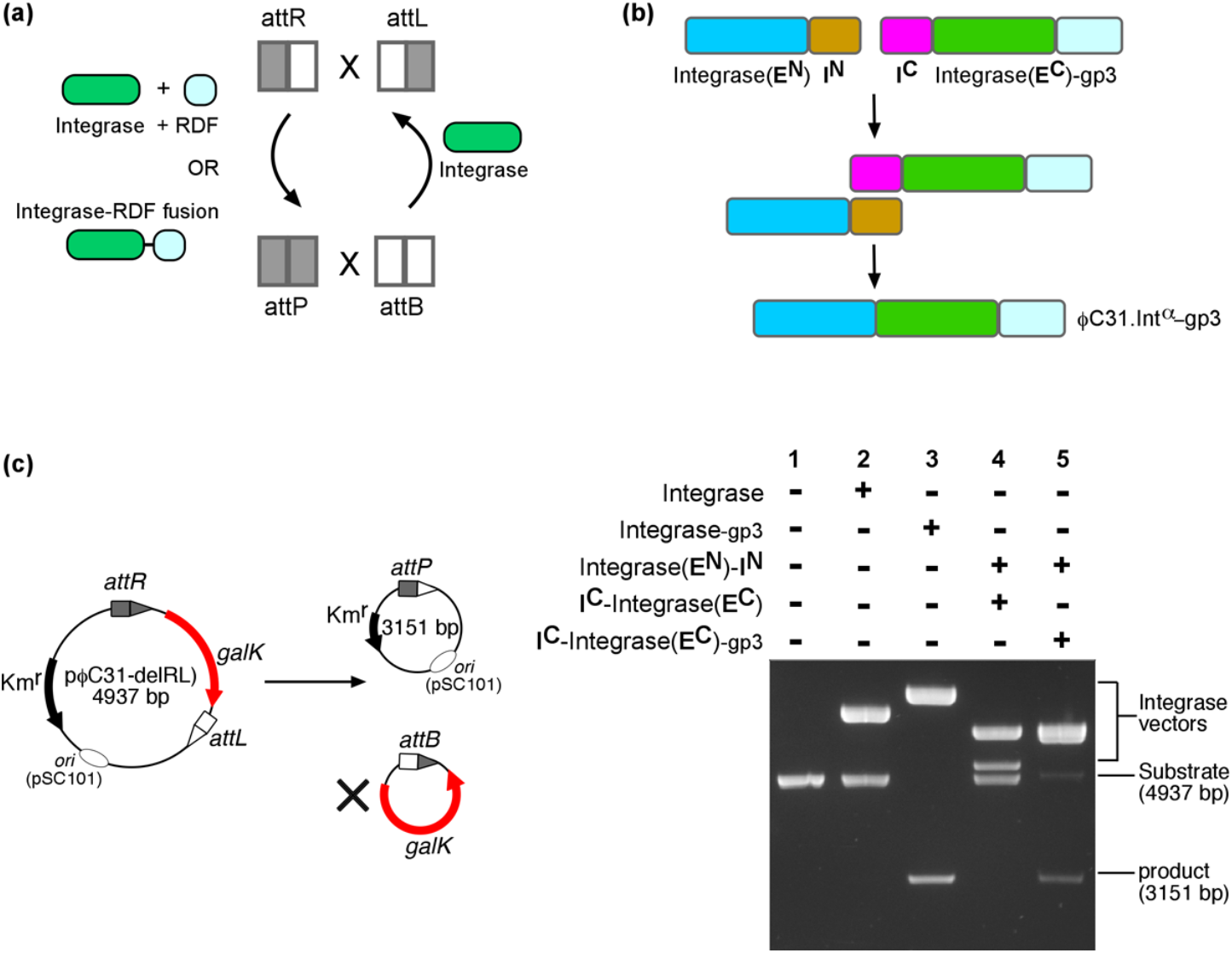
Intein-mediated assembly of *φ* C31 integrase-RDF fusion recombinase and catalysis of *attR* x *attL* recombination. **(A)** Catalysis of *attR* x *attL* recombination by integrase + RDF, or integrase-RDF fusion protein (see text for details). The RDF (Recombination Directionality Factor) for *φ*C31 integrase is gp3. **(B)** Split-intein catalyzed assembly of integrase-RDF fusion recombinase. Non-covalent association of Integrase-**E**^n^-**I^N^** and **I**^C^-Integrase-**E**^C^-gp3 via the intein domains (**I**^N^ and **I**^C^) is followed by a trans-splicing reaction that gives the fusion recombinase *φ*C31.Int^α^-gp3. **(C)** Assay of recombination activities *(galK* colour assay) of integrase constructs on *attR × attL* deletion substrates (p*φ*C31-delRL). The test substrate is identical to p*φ*C31-delPB (Figure 3A), except that the *attP* and *attB* sites are replaced by *attR* and *attL* respectively. The *in vivo* recombination assay and analysis are as described in Figure 4C. (1) Substrate only (blank control); (2) FEM141: *φ*C31.Integrase; (3) FEM33: *φ*C31.Integrase-gp3 fusion; (4) FEM136 (Integrase-**E^N^**-**I^N^**) and FEM137 (**I**^C^-Integrase-**E**^C^); (5) FEM136 (Integrase-**E^N^**-**I^N^**) and FEM188 (**I**^C^-Integrase-**E**^C^-gp3).

To demonstrate the use of our split-intein regulated system in *attR* x *attL* recombination, we split *φ*C31.integrase-gp3 between the β9 and αI domains of the integrase (Figure 5B), at the same position as described above for *φ*C31 integrase itself. To test for recombination activity, we used a test substrate (p*φ*C31-delRL) similar to the *attP* x *attB* substrate (p*φ*C31-delPB) but with the *attP* and *attB* sites replaced by *attR* and *attL* respectively (Supporting Information, Table S1). The sites are arranged in the head-to-tail orientation such that deletion of the *galK* gene leads to a reaction product that is smaller in size than the starting substrate (Figure 5C). As expected, neither integrase (Figure 5D, lane 2) nor co-expression of the Integrase-**E^N^**-**I^N^** and **I**^C^-Integrase-**E**^C^ (lane 4) were able to catalyse *attR* x *attL* recombination. Co-expression of the intein-tagged integrase N-extein (Integrase-**E^N^**-**I^N^**) and C-extein-gp3 (**I**^C^-Integrase-**E**^C^-gp3) results in efficient *attR* x *attL* recombination (lane 5). The activity of the *trans*-spliced fusion recombinase (*φ*C31.Int^α^-gp3) was comparable to that observed from native *φ*C31.integrase-gp3 (lane 3).

These experiments demonstrate that our split-intein regulated recombination system can be used to toggle between two states in which the forward reaction is catalyzed by the serine integrase, and the reverse reaction by the integrase-RDF fusion. In a practical application, reconstitution of the integrase and the integrase-RDF fusion could be regulated by two pairs of orthogonal split inteins. Several split inteins have been described in the literature^43,44^. Further characterization of their properties would make them available as orthogonal functional parts for regulation of serine integrase-based applications. More recently, mutations at residues within the GEP loop of the *Npu* DnaE split intein have been identified that affect the specificity, reaction rates, and tolerance to changes in the flanking residues^16,17^. Use of these intein variants with broader specificities would require minimum changes to the extein sequence of the target integrase, thereby reducing the potential risk of introducing deleterious mutations. Such promiscuous split inteins that have more flexible requirements for flanking residues should make it easier to find new split positions in a wider range of target proteins.

### Conclusion

Serine integrases promote efficient directional DNA site-specific recombination, and thus have major potential applications in genome engineering and metabolic pathway engineering^9,11,45–48^. They have also been incorporated into the design of cellular state machines and biocomputing devices^4,8,11^. Here we have demonstrated a novel method for regulation of serine integrase activity by making activity dependent on post-translational *trans*-splicing of two integrase extein components using split-intein technology. Since *in vitro* protein splicing by some split inteins has very fast kinetics^24,44^, this technology can be developed for the expression and purification of serine integrases that are toxic to the expression host^12^. This could significantly increase the range of integrase specificities available for *in vitro* applications such as genome engineering and DNA fragment assembly.

## MATERIALS AND METHODS

### Plasmids and DNA

The DNA sequences of all protein constructs and plasmid substrates used in this study are shown in Supporting Information (Table S2). The codon-optimized *φ*C31 integrase sequence was derived from pFM141^30^. Codon-optimized sequences of NpuN, a 102-amino acid residue split intein from *Nostoc punctiforme* DnaE, and SspC, a 36-amino acid residue split intein from *Synechocystis sp.* DnaE, were from GeneArt (Invitrogen). Plasmids for constitutive expression of the N-terminal extein-intein fusion protein (Integrase-Ext^N^-**I^N^**) in *E. coli* were made by inserting protein-coding DNA sequences between NdeI and Acc65I sites in pMS140, a low-level expression vector with a pMB1 origin of replication^30^. Plasmids for constitutive expression of the C-terminal intein-extein fusion protein (**I**^C^-Integrase-**E**^C^) were made in a similar way by inserting the coding sequence between NdeI and Acc65I sites in pEK76, which has a p15a (pACYC184) origin^30,49^. Plasmids for tet-inducible expression of the split integrase components were made by cloning the protein-coding DNA sequences between SacI and SalI sites in pTet^50^ while those for ara-inducible expression were made by cloning the protein-coding sequences between SacI and Sal sites in pBAD^51^.

The plasmid substrate for assessing recombination (deletion) by native *φ*C31 integrase or *trans*-spliced *φ*C31 integrase (p*φ*C31-delPB) was described in Olorunniji *et al.^30^*. The test substrate contains a *galK* gene flanked by *attP* and *attB* sites arranged in direct repeat (head-to-tail orientation) resulting in deletion of the *galK* gene upon integrase-catalysed recombination. The plasmid substrate for assessing recombination (inversion) activity (p*φ*C31-invPB) was made by cloning the invertible promoter device shown in Figure 3A in a pSC101 origin plasmid with kanamycin resistance selection.

Full sequences of all plasmids are shown in Supporting Information (Table S2).

### Recombination analysis

Assays for integrase-mediated recombination were carried out in *E. coli* strain DS941^30^. DS941 was first transformed with the test substrate plasmid (p*φ*C31-delPB or p*φ*C31-invPB). The substrate plasmid-containing strains were then either transformed with an integrase-expressing plasmid (pFM141), or co-transformed with two plasmids, each expressing part of the split integrase. The transformants were grown on selective MacConkey-galactose indicator plates (MacConkey agar base (Difco) supplemented with 1% galactose); kanamycin (50 μg/ml) was included to select for the substrate plasmid and the recombination product, while ampicillin (50 μg/ml) and/or chloramphenicol (25 μg/ml) were included to select for the integrase expression plasmids. Integrase-mediated deletion of the *galK* gene results in pale-coloured (*galK*^-^) colonies on the indicator plates, whereas red (*galK*^+^) colonies suggest lack of recombination. In some cases, colonies in which low-level recombination has occurred maintain a red colour. To determine the extent of recombination more accurately, plasmid DNA recovered from the cells was examined. L-broth (1 ml) was added to the surface of each plate and the cells were suspended using a plate spreader. An aliquot of this suspension was used to inoculate L-broth (1 in 1000 dilution), and the culture was incubated overnight at 37 °C with kanamycin selection for the substrate and recombinant plasmids. Plasmid DNA was prepared using a Qiagen miniprep kit, and analysed by 1.2% agarose gel electrophoresis.

### Inducible expression of integrase-intein fusions for *trans-splicing*

*In vivo* expression of the integrase-intein fusion proteins under the control of arabinose and tet inducible promoters was carried out in strain DS941/Z1 (S.D. Colloms, unpublished), which consitutively expresses TetR, required for regulation of the pTet promoter. The DS941/Z1 strain was made competent by a standard calcium chloride method^52^. The cells were transformed with the plasmid substrate p*φ*C31-invPB, then cultured for 90 minutes and selected on L-agar plates containing kanamycin (50 μg/ml). A single colony was picked and grown in kanamycin-containing L-broth (5 ml) to make a stationary phase overnight culture. An aliquot of the stationary phase culture was then diluted into L-broth containing kanamycin and grown to mid-log phase. These cells were made ‘chemically competent’ (as above) and transformed with the two vectors containing the coding sequences of the intein-integrase fragments. The plasmid vector pFEM148 (ampicillin selection) expresses Integrase-**E^N^**-**I^N^** under the control of the pTet promoter. The second plasmid pFEM149 (chloramphenicol selection) expresses **I**^C^-Integrase-**E**^C^ under the control of the pBAD promoter.

The transformant cells were cultured for 90 minutes, and selected on L-agar plates containing kanamycin (50 μg/ml), ampicillin (100 μg/ml), and chloramphenicol (25 μg/ml). To carry out the recombination assays, a culture from a single colony was grown overnight in L-broth in the presence of the three antibiotics, to stationary phase. The culture was diluted further (1:100) and grown to mid-log phase (about 90 minutes), after which expression of the split-intein-integrase fragments was induced by the addition of anhydrotetracycline, aTc (0.1 μg/ml), arabinose, Ara (0.2% w/v), or both, as shown in Figure 3C. Glucose was added to 0.4% concentration to the cultures where it is required to turn off expression of the arabinose promoter. The induced cultures were grown for 24 hours at 37 °C, after which the cultures were left at room temperature for 8 hours. Next, aliquots of each culture (50 μl) were diluted with 950 μl phage buffer (10 mM Tris, pH 7.5, 10 mM MgCl_2_, 68 mM NaCl), and GFP expression was measured by means of fluorescence.

### Fluorescence measurements

Fluorescence measurements were carried out on a Typhoon FLA 9500 fluorimager (GE Healthcare). Aliquots of the diluted cultures (200 μl) were added to a 96-well plate, and the fluorescence of the expressed proteins were measured (GFP: excitation, 485 nm; emission, 520 nm and RFP: excitation, 532 nm; emission, 575 nm). To determine cell density for each sample, 50 μl aliquots were diluted to 1000 μl, and the spectrophotometric absorbance was read at 600 nm. The GFP-independent background signals of the cells were determined by measuring the fluorescence of strain DS941/Z1 strain (containing the test substrate but without the split-intein-integrase expression vectors). The background fluorescence was subtracted from the values measured for samples in the different treatment groups, after normalization using the cell density measurements.

## SUPPORTING INFORMATION

Table S1: Sequences of *φ*C31 integrase *att* sites.

Figure S1: Comparison of *in vivo* recombination activity of native *φ*C31 integrase with the reconstituted *trans*-spliced *φ*C31 integrase.

## ACKNOWLEDGEMENTS

We thank Dr Sean Colloms and Dr Steve Kane for the gift of the plasmid template (pSWITCH) used in constructing the invertible promoter assay substrate.

## SUPPORTING INFORMATION

**Figure S1:**
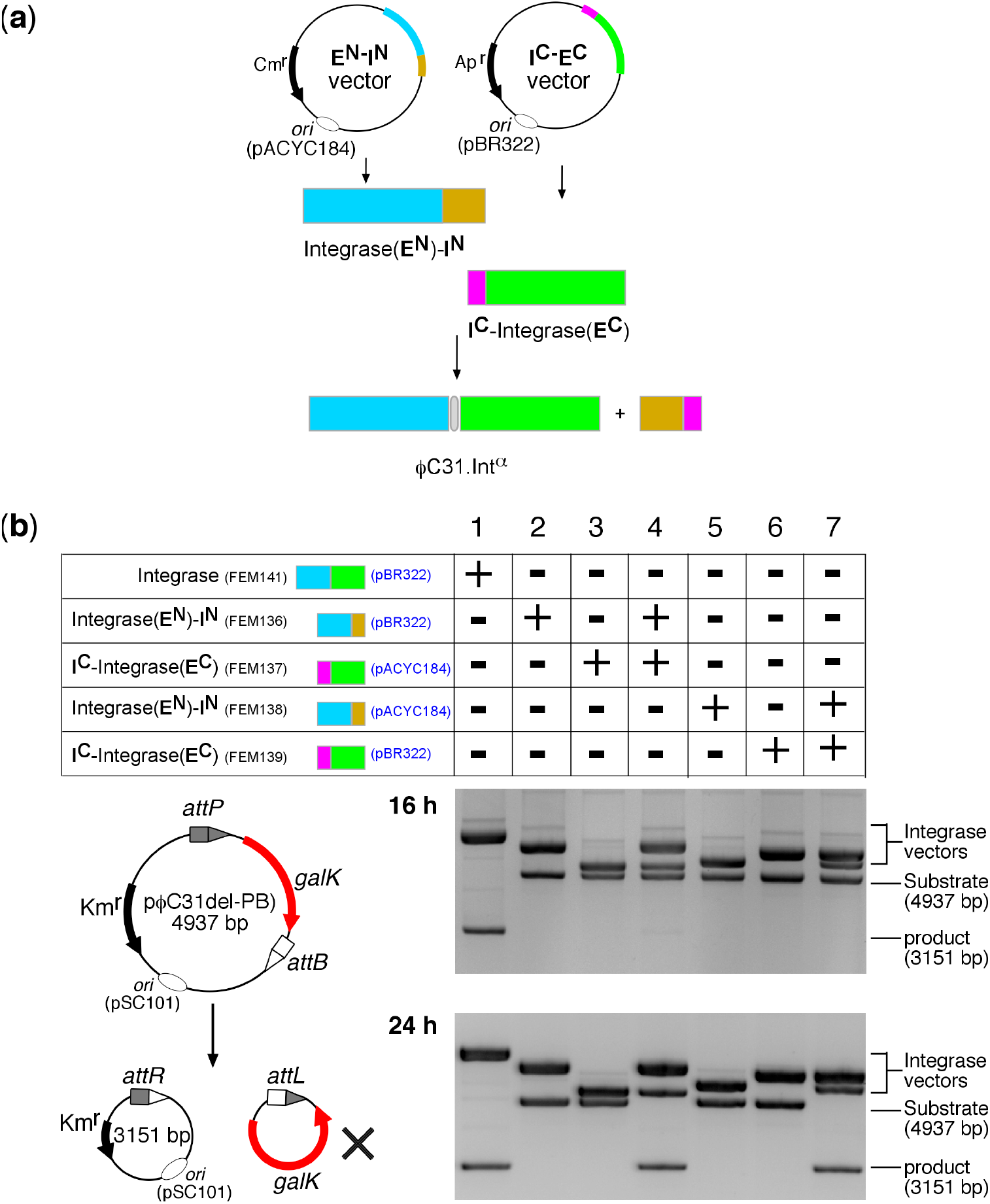
Comparison of *in vivo* recombination activity of native *φ*C31 integrase with reconstituted trans-spliced *φ* C31 integrase. **(A)** The precursor polypeptides for the split-intein integrase were consitutively expressed from vectors with either a pACYC184 or pBR322 origin of replication (Proudfoot *et al.,* 2011). For each precursor, both types of vector were tested, to confirm that the expression vector did not have any serious effect on the outcome of the experiment (see part B). In the example shown, the **E^N^**-**I^N^** vector is expressed in pACY184, and the **I**^C^-**E**^C^ construct placed in pBR322 vector. **(B)** Assay of recombination activities *(galK* deletion assay) of integrase constructs on an *attP × attB* substrate (p*φ*C31-delPB). The *in vivo* recombination assay and analysis are as described in Figure 4C. Cells containing p*φ*C31-delPB were transformed with the expression vectors indicated and grown for 16 hours (top panel, **16 h**) or for 24 hours (bottom panel, **24 h**) on selective plates. For analysis of *in vivo* recombination products, plasmid DNA was recovered from cells (Olorunniji *et al.,* 2017) and separated by means of 1.2% agarose gel electrophoresis.

**Table S1.**
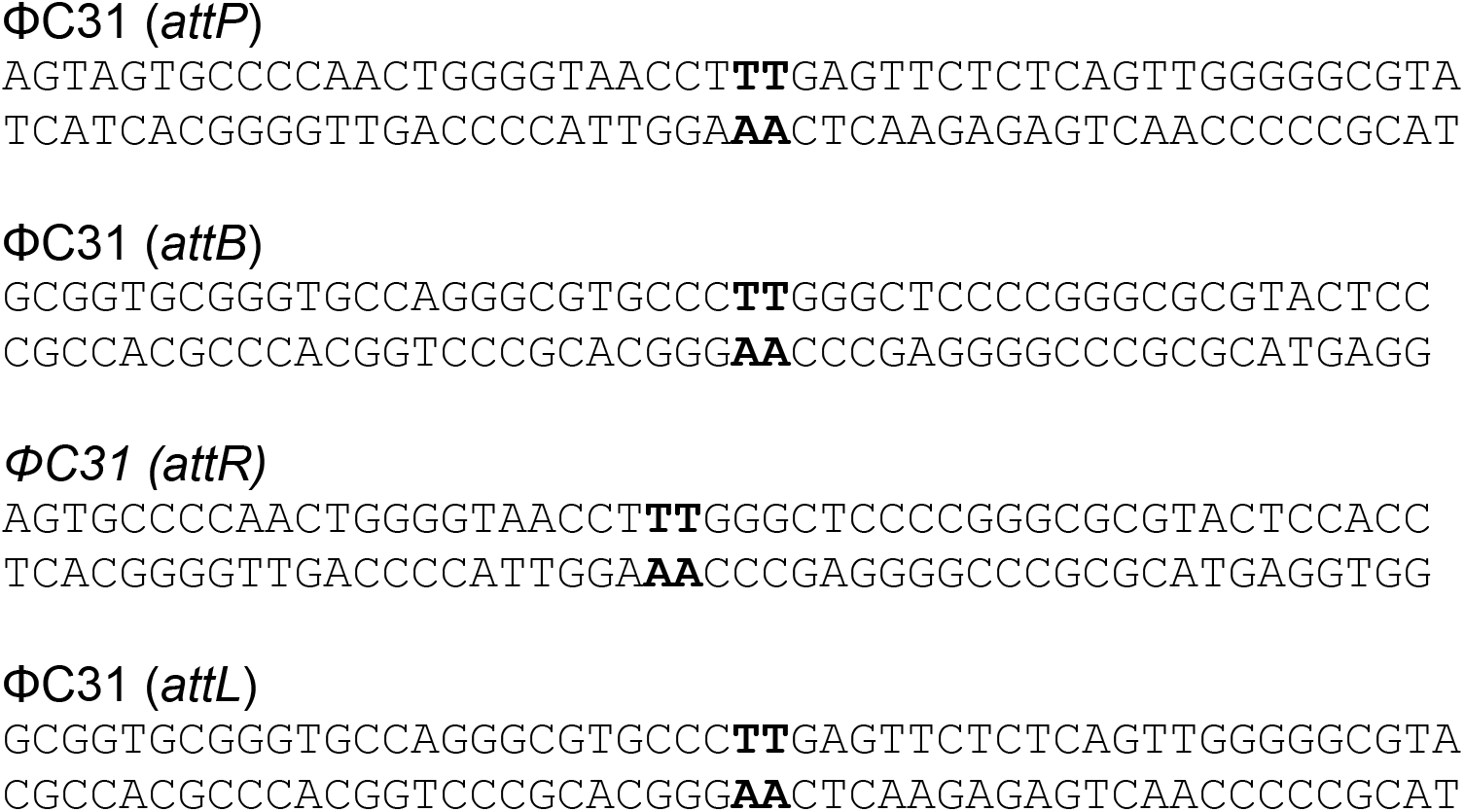
Sequences of ΦC31 integrase att sites. The sequences are shown in the head-to-tail orientation that results in deletion of the intervening sequences upon recombination. The central 2-bp overlap sequences of the att sites are highlighted in bold text.

